# Colonization of *Bacillus altitudinis* on the Compatible Soybean Varieties to Provide Seed Rot Resistance

**DOI:** 10.1101/2023.11.27.568843

**Authors:** Ping-Hu Wu, Hao-Xun Chang

## Abstract

Seed health is crucial for plant growth and agricultural productivity. Recent studies have illustrated the importance of plant microbiome in disease resistance, however, it remains unclear whether the seed microbiome confers seed rot resistance against fungal pathogens. In this study, the application of antibiotics on the seeds of eight soybean varieties showed that seed-associated bacteria were involved in the seed rot resistance caused by *Calonectria ilicicola*, but this resistance cannot be carried to withstand root rot. Using PacBio 16S rDNA full-length sequencing and microbiome analyses, the seed microbiome was shown to mainly dependent on the soybean variety, and there was no consistent community network associated with seed rot resistance across soybean varieties. Instead, the seed-associated *Bacillus altitudinis* was identified through the differential abundance analysis and culture-dependent isolation. Moreover, qPCR confirmed the persistence of *B. altitudinis* on apical shoots till 21 days post-inoculation, but not on roots by 9 days post-inoculation. The short-term colonization of *B. altitudinis* on roots may explain the absence of root rot resistance. Furthermore, seed treated with *B. altitudinis* restored seed rot resistance, but only in the compatible soybean varieties. For the incompatible soybean varieties, *B. altitudinis* showed lower bacterial density and provided no seed protection. Collectively, this study advances the insight of *B. altitudinis* conferring seed rot resistance. These findings highlight the potential of using seed-associated bacteria for seed protection and underscore the importance of considering bacterial compatibility with plant genotypes and tissues.

## Introduction

The plant microbiome constitutes a vast and complex community of microbes, playing a crucial role in supporting plant health, productivity, and resilience [1]. These microbes engage in various beneficial interactions with their hosts, such as nutrient acquisition, growth promotion [2, 3], and improving resistance to biotic and abiotic stresses [4–6]. As a result, the plant microbiome has received significant research attention due to its potential to sustain agriculture and the environment [7, 8]. While research over the past decade has predominantly concentrated on the phyllosphere and rhizosphere microbiomes, the seed microbiome, or spermosphere microbiome, has been relatively overlooked, despite its fundamental importance in plant growth and agricultural production.

The composition of the seed microbiome is shaped by factors such as plant species, genotypes, and environmental conditions. For example, domestication has been shown to decrease microbial diversity in wheat seeds, whereas domesticated rice exhibits a greater microbial diversity than their wild ancestors [9, 10]. Moreover, investigations indicate that the seed endophytic microbiome of rapeseed, pumpkin, and tomato can be influenced by host cultivar/genotype and environmental factors [11–13], while the seed epiphytic microbiome is primarily affected by environmental factors such as location [14]. It is important to note that methodological variations, including the surface sterilization procedure, can impact the detected species richness in different studies [15, 16]. Nevertheless, there is a consensus across studies that seeds harbor fewer microbial species compared to other plant tissues [17]. This reduction in diversity raises questions about how plants select compatible seed-associated microbes and their impact on plant health.

One of the initial perspectives on seed-associated microbes stemmed from the recognition of seed-borne pathogens, which can cause diseases and significantly impact seed health [18]. This understanding led to the widespread adoption of physical and chemical seed treatments, such as hot water soaking and fungicide coating, to eliminate seed-associated microbes in conventional farming. However, recent research is reshaping this perspective, unveiling a diverse range of beneficial bacteria and fungi in the spermosphere. These seed-associated microbes are now recognized for playing crucial roles in seed germination, seedling development, and seedling protection from pathogens [19]. For instance, studies have demonstrated that seeds of rice and millet treated with antibiotics, resulting in the absence of bacteria, exhibited slower germination processes [20, 21]. Similarly, seeds of maize and pearl millet treated with antibiotics showed reduced seedling growth [22, 23]. Furthermore, endophytic bacteria such as *Bacillus*, *Pseudomonas*, and *Sphingomonas* have been identified as contributors to protect young seedling through the production of antimicrobial substances [5, 24] or the stimulation of plant defense responses [25, 26]. Some of these microbes can be vertically transmitted from parents to offspring plants, ensuring the continuity of a beneficial holobiont across generations [11, 17, 27]. Accordingly, these findings suggest that the seed microbiome can be considered a heritable and extended phenotype. This concept not only reshapes the traditional approach of seed sterilization for disease management [28] but also underscores the potential of harnessing the seed-associated microbes to develop sustainable seed protection strategies.

Soybean (*Glycine max*) holds significant agricultural importance globally. One of the primary challenges in soybean cultivation is the prevalence of seed-borne and soil-borne diseases, which can hinder seed germination through seed rot, root rot, and damping-off [29]. In the United States, these diseases accounted for 76% of soybean yield loss related to diseases from 2018 to 2020 [30]. Various fungal pathogens like *Athelia rolfsii*, *Fusarium oxysporum*, *Macrophomina phaseolina*, *Rhizoctonia solani*, and *Sclertoinia sclerotiorum* have become persistent endemic problems [31], and a recently emerging disease, red crown rot (RCR) caused by *Calonectria ilicicola*, has gained attention globally in recent years [32]. While fungicide treatments are commonly suggested to manage both endemic and emerging diseases [31, 33], alternative strategies such as plant resistance and beneficial microbes offer potentially more sustainable options for disease management. Considering the potential demonstrated by the seed microbiome of other plants in managing these diseases, exploring the soybean seed microbiome for disease management becomes a viable option. Current studies on the soybean seed microbiome indicated lower diversity compared to rhizosphere and phyllosphere microbiomes [34]. Using culture-dependent and culture-independent methods, 22 and 27 bacterial genera were identified in the soybean seed microbiome, respectively [35, 36]. Among these seed-associated microbes, *Bacillus*, *Pantoea*, and *Sphingomonas* were identified as the most dominant genera in soybean seeds [36], and some of these seed-associated bacteria exhibited *in vitro* antagonistic activity against various soybean pathogens [35].

While previous investigations into the soybean seed microbiome relied on the variable regions of 16S rDNA, primarily V3-V4, for taxonomic profiling, a growing body of evidence underscores the advantages of employing 16S rDNA full-length sequencing. This approach offers more precise insights into microbial composition [37–39]. Despite these advantages, full-length sequencing has not been widely utilized in studying the soybean seed microbiome. In our research, we utilized 16S full-length sequencing to explore the seed microbiome of diverse soybean varieties, differing in seed rot resistance but not root rot resistance to *C. ilicicola*. Our findings underscored that the discreteness between seed rot resistance and root rot resistance could be attributed to the seed-associated bacteria. Furthermore, we verified that the colonization of *Bacillus altitudinis* on the seeds of compatible soybean varieties is crucial for preventing seed rot. This study presents the first instance of seed-associated *B. altitudinis*, exhibiting not only antagonistic effects against various soil-borne fungal pathogens, but also contributing to seed rot resistance through seed colonization.

## Material Methods

### Preparation of fungal and plant materials

Fungal materials including *Athelia rolfsii* isolate a31, *Calonectria ilicicola* isolate F018, *Fusarium oxysporum* isolate R1031, *Macrophomina phaseolina* isolate 1-4-03, *Rhizoctonia solani* AG-7 isolate WDG070, *Sclerotinia sclerotiorum* isolate 1980, were routinely cultured on the potato dextrose agar (PDA) plates (HIMEDIA, India), and maintained on filter papers at -20 °C or preserved in 20% glycerol at -80 °C for long-term storage.

Upon experimental setup, soybean seeds were triply washed with tap water and then immersed in 1% NaOCl for 10 min, followed by repeatedly rinsing with sterile distilled H_2_O for five times. The completeness of surface sterilization was assessed by spreading 100 µl aliquot of the last rinse on the nutrient agar (NA) plates (HIMEDIA), and then incubated at 28°C for 5 days. Seeds were considered surface-disinfected and would be used in subsequent experiments if no microbial colony was formed on the NA plates. There are 16 varieties of soybean seeds used in this study (Table S1).

### Phenotyping soybean seeds in response to the inoculation of *Calonectria ilicicola*

The seed rot assay was conducted according to Broders et al. [40] with slight modifications. In summary, conidia were collected from a ten-day-old *C. ilicicola* colony on ½-strength PDA (½PDA) and diluted to two conidia per microliter. 100 µL of the diluted conidia (200 conidia) were spread on the 1.5% water agar (WA) plates and incubated at 25°C in dark for 2 days. Subsequently, eight surface-disinfected soybean seeds were placed on each WA plate and cultured at 25°C in dark for 5 days. Surface-disinfected soybean seeds of each variety on the WA plates without conidia were included as the controls. Soybean seeds were considered dead for situations including no germination, the radicle length was less than 1 cm, or the cotyledon and radicle were fully colonized by mycelia. The seed rot resistance was represented by the mortality rate, meaning the number of dead seeds divided by the eight seeds in each WA plate. Each plate was counted as a biological replicate, and each experiment was constituted of three biological replicates. The experiment was repeated three times independently.

For the cotyledon rot assay, the severity of cotyledon rot was scored on a six-grade scale based on the lesion size and severity on the cotyledon: 0 = no lesion; 1 = lesion less than 25 %; 2 = lesion less ranging from 26 to 50 %; 3 = lesion less ranging from 51 to 75 %; 4 = lesion over 75 %; 5 = seeds with no germinate or all cotyledon and radicle were colonized by mycelia. Finally, the seed rot severity was calculated by standardizing the seed mortality and the cotyledon rot to range from 0 to 1 and averaging these two indices.

For the root rot assay, the fungal inoculum was freshly prepared according to the previous methods [33], and 15 mL of the fungal inoculum or control inoculum was mixed with commercial potting soils (T033, Garden Castle Ltd., Taiwan) in 500 ml pots. Each pot contained four soybean seeds, and these pots were placed in a greenhouse at 25 °C in a 16h-8h light-dark cycle with daily irrigation. After 21 days, soybean seedlings were collected to separate roots from soils by gentle rinsing with tape water. The disease severity was visually scored using a six-grade scale: 0 = no symptom; 1 = small brown necrotic lesions on the primary root; 2 = brown necrotic lesions extending over the primary root and some lateral roots; 3 = over half root lost and rotted with brown necrosis on the subterranean stem; 4 = almost all roots lost and rotted; and 5 = dead seedlings [41]. Each treatment included three biological replicates (pots) in a complete randomized design, and the experiment was repeated three times independently.

### Elimination of seed-associated bacteria by antibiotics

To assess the role of seed-associated bacteria in seed rot resistance, soybean seeds were firstly surface-disinfested as abovementioned method, and then soaked in a solution containing ampicillin (100 µg/mL), rifampicin (100 µg/mL), and streptomycin (100 µg/mL) for 16 h (Fig. S1). The control group was soaked in sterile distilled H_2_O for the same period. Following treatment, the seeds were rinsed 7 times with sterile distilled H_2_O to remove antibiotics residue. To ensure the completeness of eliminating seed-associated bacteria, the seeds were ground using a mortar and pestle in 2-fold volume (v/w) of phosphate-buffered saline (PBS) buffer, and 100 µL of the grinding aliquot was spread onto the tryptic soy agar (TSA) plates (STBIO MEDIA). After incubating the TSA plates at 28°C for 5 days, the elimination of seed-associated bacteria was considered successful if no colony was observed.

### PacBio 16S rDNA full-length sequencing and analyses of the seed microbiome

DNA was extracted from 5 day-post-germination soybean seeds using the CTAB method. The bacterial 16S rDNA region was amplified by the universal primers 27f and 1492r (Table S2), sourced from the PacBio library preparation kit. To minimize the amplification of host DNA, the PNA blocker targeting soybean chloroplast DNA [42] was incorporated at a final concentration of 2.5 pmole, and the LNA blockers were added to target soybean mitochondria DNA [43] at a final concentration of 5 pmole (2.5 pmole for each direction). The sequencing libraries were prepared according to the workflow of the PacBio SMRTbell kit, and sequencing was performed on the PacBio Sequel IIe platform with 10 h movie collection time.

The raw sequencing files were filtered, trimmed, and de-replicated using the PacBio single-molecule real-time link software to generate circular consensus sequencing reads. The R package “DADA2” v1.26 was utilized to denoise and construct amplicon sequence variants (ASVs) following the adjusted parameters [44, 45]. The chimeras were removed before classifying the ASVs using the Bayesian classifier in DADA2 and the SILVA v132 database at 99% similarity [46]. The non-prokaryotic and unclassified ASVs were removed before finalizing the ASV table. To assign the taxonomy to each ASV, the sequences were aligned using BLAST+ [47] to NCBI 16S rRNA gene database with an E-value at 10^-5^. The classification of each ASV was determined by the lowest E-value, followed by the highest Bit score, and then the highest identity. The BLAST results were processed by the R package “taxize” v0.9.1, resulting in a final taxonomy table. The ASV sequences were aligned using MAFFT [48], and a maximum likelihood phylogenetic tree was constructed with IQ-TREE2 with 1,000 bootstrap replicates [49].

The ASV table, taxonomy table, and phylogenetic results were imported into the R package “phyloseq” v1.42, and four samples with fewer than 2,000 reads were filtered [50]. ASVs with a mean of relative abundance below 0.01% across 36 samples or with occupancy below 5% (lower than 2 samples) were excluded from the subsequent analyses. To assess α-diversity indices, all samples were rarefied to the sample with the lowest sequencing depth using the R package “vegan” v2.64-4 and “picante” v1.8.2 [51, 52]. To assess the β-diversity, ASVs were normalized by median sequencing depth before subjected to non-metric multidimensional scaling (NMDS) based on the Bray-Curtis distance, and three NMDS axes were included to obtain smaller stress. The cluster analysis was calculated by the beta flexible method (with α = 0.625) performed by “agnes” function in the R package “cluster” version 2.1.4 using the Sørensen distance. PERMANOVA was conducted using the “adonis2” function in “vegan” package with 999 permutations. The differential abundance of ASVs between the resistant and susceptible varieties was analyzed using the R package “DESeq2” v1.38.3 [53], and the significance was determined at the BH-adjusted *p*-values at 0.05. For the network analysis, the co-occurrence of ASVs in each soybean variety was calculated using ‘sparcc’ algorism provided in the R package “NetComi” v1.1. Correlations larger than 0.7 with *p*-values less than 0.05 were kept to construct the network topology by NetComi. The hub taxa were identified based on the eigenvector centrality values above the 95% quantile of a fitted log-normal distribution [54].

### Isolation of seed-associated bacteria and *in vitro* antagonistic assay against fungal pathogens

Surface-sterilized soybean seeds were germinated on the 1.5% WA plates for 5 days before grinding using a mortar and a pestle. The ground aliquots were serial diluted till 10^-5^ folds using PBS buffer, and the 100 µL diluted aliquot was spread on the soymilk agar (SA) plates, NA plates, Luria-Bertani agar (LA) plates, and 0.1% TSA plates. The plates were incubated at 28 °C for 7 days, with three plates for each dilution fold. After incubation, colonies were differentiated and selected based on their morphological features. The selected colonies were subsequently single colony purified using the streak plate technique. Finally, the purified colonies were preserved in 25% glycerol at -80°C for long-term storage.

The seed-associated bacteria were tested for their antifungal activity against fungal pathogens in the dual culture assay. A mycelial plug (5 mm in diameter) from the actively growing edge of each fungal species was placed on the center of a medium plate. The bacteria were cultured in the tryptic soy broth (TSB) for 16 h and adjusted to OD_600_ value of 1. Subsequently, 2 µL of the bacterial aliquot was placed 3 cm from the mycelial plug on both sides of the TSA medium. This method tested fungal pathogens under varying conditions of temperature and measurement times. Specifically, *A. rolfsii* and *C. ilicicola* were measured at 7 days post-inoculation (dpi), *M. phaseolina* and *S. sclerotiorum* at 5 dpi, and *R. solani* at 2 dpi. All fungi were cultured at 28 °C in the dark, except for *S. sclerotiorum* which was cultured at 25 °C. The inhibition rate was calculated by:

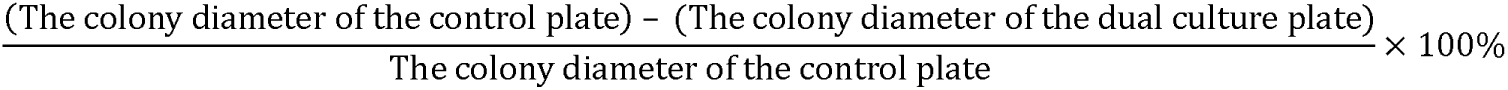

For identifying the species of TN5S8 and TN3S3, the genomic DNA were extracted using the Presto gDNA Bacteria Advanced Kit (Geneaid Biotech Ltd, Taiwan), and subjected to PCR using the 27F/1492R for 16S rRNA gene and UP1/2r for *gyrB* gene [55] (Table S2). The amplicons were submitted for Sanger sequencing (Genomics, Taiwan), and sequencing results were subjected to BLAST search in the NCBI database and used for phylogenetic analysis.

### Colonization of *Bacillus altitudinis* TN5S8 on different soybean varieties

Antibiotics-treated soybean seeds were soaked for 16 h in the freshly prepared aliquot of *B. altitudinis* TN5S8 (hereafter abbreviated as TN5S8). Meanwhile, the control group was soaked in PBS buffer. The inoculant was prepared from a 24-hour-old bacterial culture in TSB, pelletized by centrifugation at 14,000 g for 3 min, before resuspended and adjusted to a concentration of 10^7^ CFU/mL using PBS buffer. The treated seeds were air-dried in a laminar flow hood. Seed rot resistance was evaluated using the plate assay method abovementioned. The colonization efficiency of TN5S8 on each soybean variety was assessed by re-isolating TN5S8 from seeds using the method aforementioned. The colony numbers per gram of seeds were assessed by:

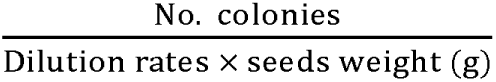

To study the impact of TN5S8 on seed germination, three different treatments were tested, including (1) surface-sterile seeds + T5S8, (2) surface-sterile seeds + cell-free culture filtrate of T5S8, and (3) antibiotics-treated seeds + T5S8. The cell-free culture filtrate was prepared from a 3-day-old bacterial culture in TSB, centrifuging at 14,000 g for 3 min and filtering the supernatant through a 0.2 µm Millex filter (Merck KGaA, Germany). The impact of T5S8 on seed germination was assessed by germination rate at 5 dpi.

### Quantitative PCR (qPCR) detection of *Bacillus altitudinis* TN5S8

The soybean TN5 seeds were surface-sterilized and inoculated with TN5S8, and then planted in pots containing a mixture of peat and perlite at a ratio of 4:1 in the greenhouse using the methods mentioned above. Plant tissue samples were collected at five timepoints. The cotyledons and epicotyls were collected for samples at 3 dpi, and the apical shoot and the first node were sampled for samples from 6 dpi onwards. The root samples were washed with sterile water to remove soil, and the taproots were collected for DNA extraction. Each biological replicate consisted of tissues from four plants, with five biological replicates obtained for each time point.

About 350 mg of plant tissues were homogenized in liquid nitrogen with a mortar and postal, and DNA was extracted using the CTAB method. Specific primers were designed to amplify a 106-bp fragment of the *gyrB* gene of *B. altitudinis* (Table S2). The qPCR was performed on the CFX Connect Real-Time System (Bio-Rad Hercules, CA, USA) using genomic DNA, iQ SYBR green supermix kit (Bio-Rad), and 0.4 μM of each primer under the following thermocycling conditions: 95 °C for 3 min; 40 cycles of 95 °C for 10 s and 57 °C for 30 s, with a melting curve processing from 60 °C to 95 °C for quality control. Soybean actin gene Glyma.15G050200 was used as an internal control to verify DNA quality. Each biological replicate was technically repeated twice. Genomic DNA of TN5S8 was serially diluted ten-fold in three separate series to obtain standards from 2 to 2.44 × 10^6^ copies per μL. Standard curves were generated by plotting the mean Ct value of the three replicates against the log-transformed DNA concentration.

### Statistical analysis

All statistical analyses were conducted using the R environment 4.2.3. For the data analyses using the *t*-test, ANOVA, and Tukey’s HSD test, homoscedasticity was checked by the Levene’s test, and normality was checked by the Shapiro test and Q-Q plot. For the data not fitting parametric assumptions, the Kruskal-Wallis test and Dunn’s test was applied. The *p* values of the Tukey’s HSD test and Dunn’s test were adjusted by the BH method for multiple comparisons. The significance of the statistical analysis was determined by α at 0.05.

## Result

### Phenotyping soybean seeds in response to the inoculation of *Calonectria ilicicola*

In assessing soybean resistance to *C. ilicicol*a, significant differences were observed for both seed mortality (*p* < 0.001) and cotyledon rot (*p* < 0.001) across 16 varieties. Soybean variety SS exhibited the highest levels of seed mortality and cotyledon rot, whereas TN5 displayed the lowest seed mortality and TN11 displayed the lowest cotyledon rot (Table S3). By averaging the seed mortality and the cotyledon rot to obtain the seed rot severity, the results showed that TN11, HBS, TN3, and TN5 were the most resistant varieties, while SS, KS9, KS7, and HC were identified as the most susceptible varieties (Fig. 1).

**Fig. 1.**
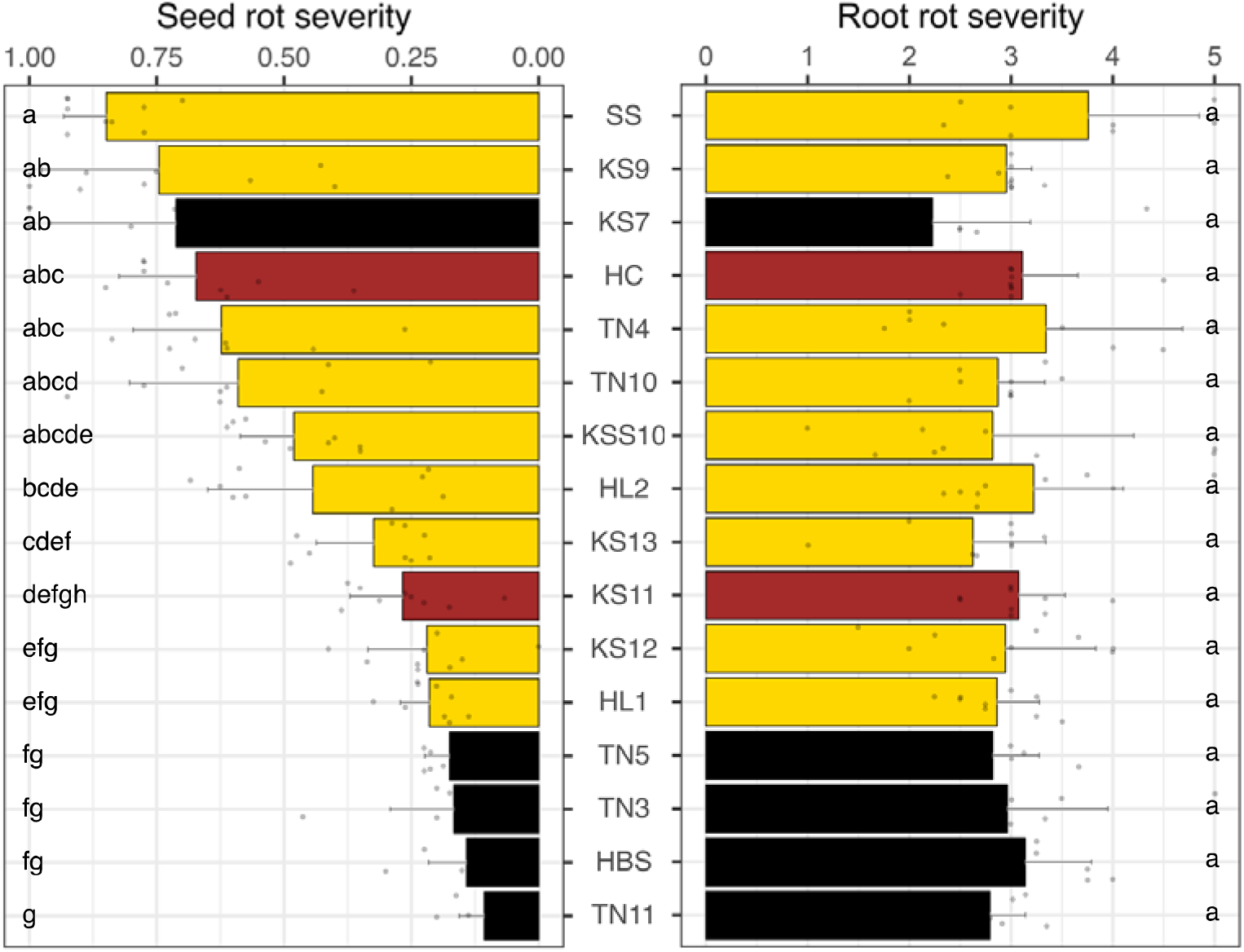
The seed rot resistance and root rot resistance of 16 soybean varieties in responses to the inoculation of *Calonectria ilicicola*. The colors indicate the color of the seed coat of the soybean variety. Values represent the means[± standard deviation. The Kruskal-Wallis and the Dunn’s test were used to determine significant difference at α = 0.05.

In contrast, no significant difference (*p* = 0.293) was observed in root rot resistance across the 16 varieties (Fig. 1). These findings suggested that resistance to *C. ilicicola* in these soybean varieties is only evident during the seed stage (seed rot and cotyledon rot) but not during the seedling stage (root rot). In other words, the discrete resistance may not be solely determined by soybean genotypes, prompting the hypothesis that seed-associated bacteria may play a role in seed rot resistance to *C. ilicicola*.

Using antibiotics-treated seeds of both resistant and susceptible soybean varieties, the susceptibility of the four originally susceptible varieties remained unchanged (Fig. 2A), but the initially resistant varieties became susceptible (Fig. 2B, C, D). As the antibiotic treatment did not affect seed germination in the control groups and did not impact the growth of *C. ilicicola*, the increased seed mortality in these four initially resistant varieties (TN11, HBS, TN3, and TN5) may be attributed to the elimination of the seed-associated bacteria.

**Fig. 2.**
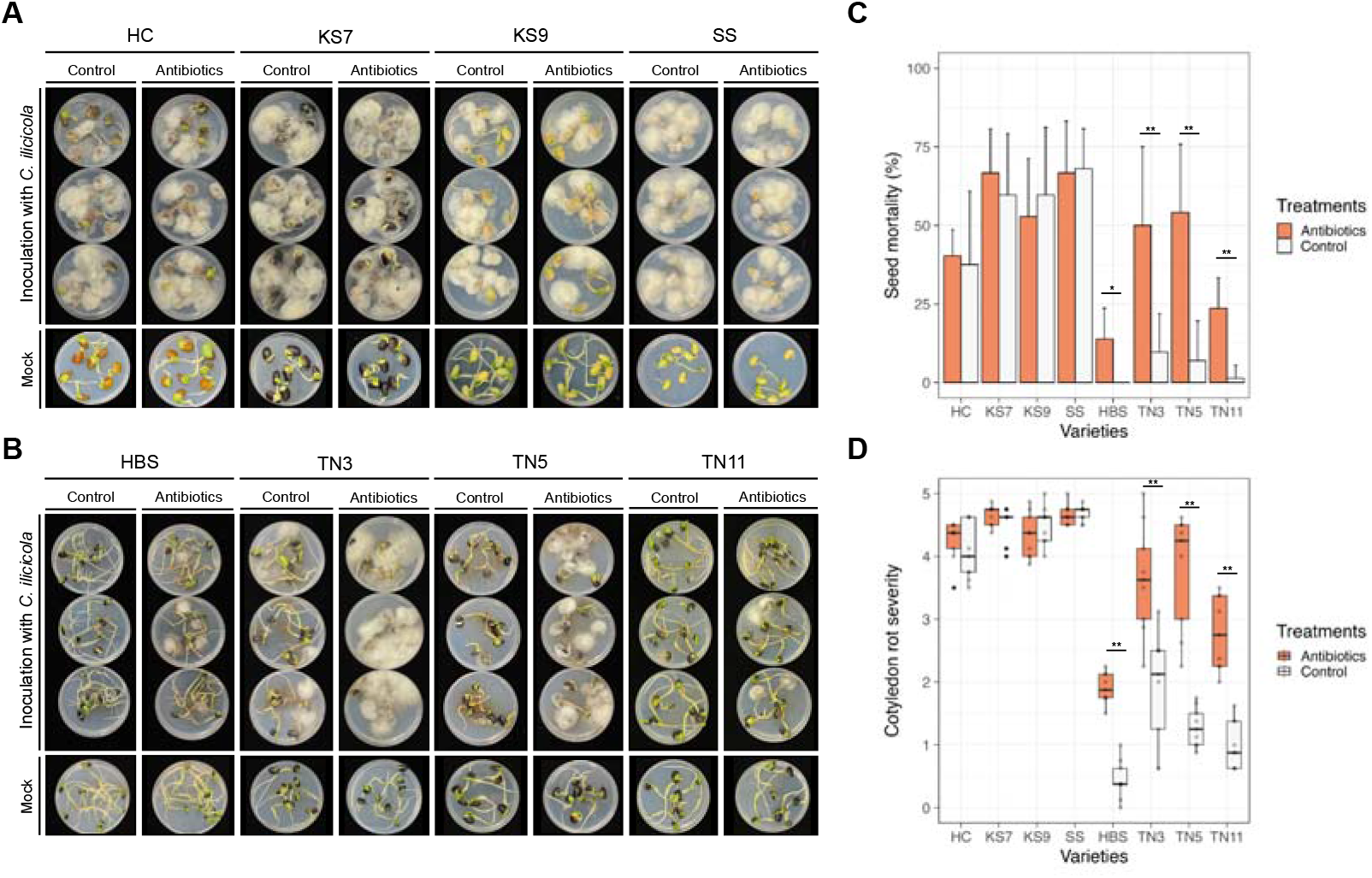
Seed rot assay using the antibiotics-treated seeds. The antibiotics included ampicillin, rifampicin, and streptomycin. The control was treated with ddH_2_O. **A** The four susceptible soybean varieties showed no difference between the control and antibiotic treatment. **B** The four resistant soybean varieties showed significant difference in seed rot between the control and antibiotic treatment. Both seed mortality and cotyledon rot severity increased for the antibiotics-treated seeds, and the four initially resistant varieties became susceptible. **C** Seed mortality. **D** Cotyledon rot severity. The orange bars represent the antibiotics-treated samples, and the white bars represent the control samples. The values are presented as means ± standard deviation. The asterisks indicate significance based on the *t*-test (*: *p* < 0.01, **: *p* < 0.001).

### PacBio 16S rDNA full-length sequencing and analyses of the seed microbiome

To delve into the seed microbiome of soybean varieties, a total of 802,727 PacBio 16S rDNA full-length raw reads were acquired. Following quality controls and the exclusion of chloroplast, mitochondrial, and non-characterized sequences, 589,009 reads were retained (Table S4). Despite the inclusion of PCR blockers, two varieties (KS7 and KS9) still exhibited interference from eukaryotic DNA (Fig. 3A). Consequently, two samples with fewer than 2,000 reads from each of the KS7 and KS9 varieties were excluded from subsequent analysis. For the remaining 36 samples, rarefaction curves indicated satisfactory sampling depth, as all curves reached saturation status (Fig. S2). After quality control, 135 ASVs were identified in the 36 samples, with BLAST results showing an identity range of 85.22% to 100% for these ASVs according to the NCBI reference taxa. Among them, 107 ASVs displayed identities greater than 99% to the reference taxa, with only one ASV having an identity below 97% (Fig. 3B). In summary, the microbiota obtained from PacBio 16S rDNA full-length sequences yielded a high-quality taxonomic profile at the species level for downstream analyses.

**Fig. 3.**
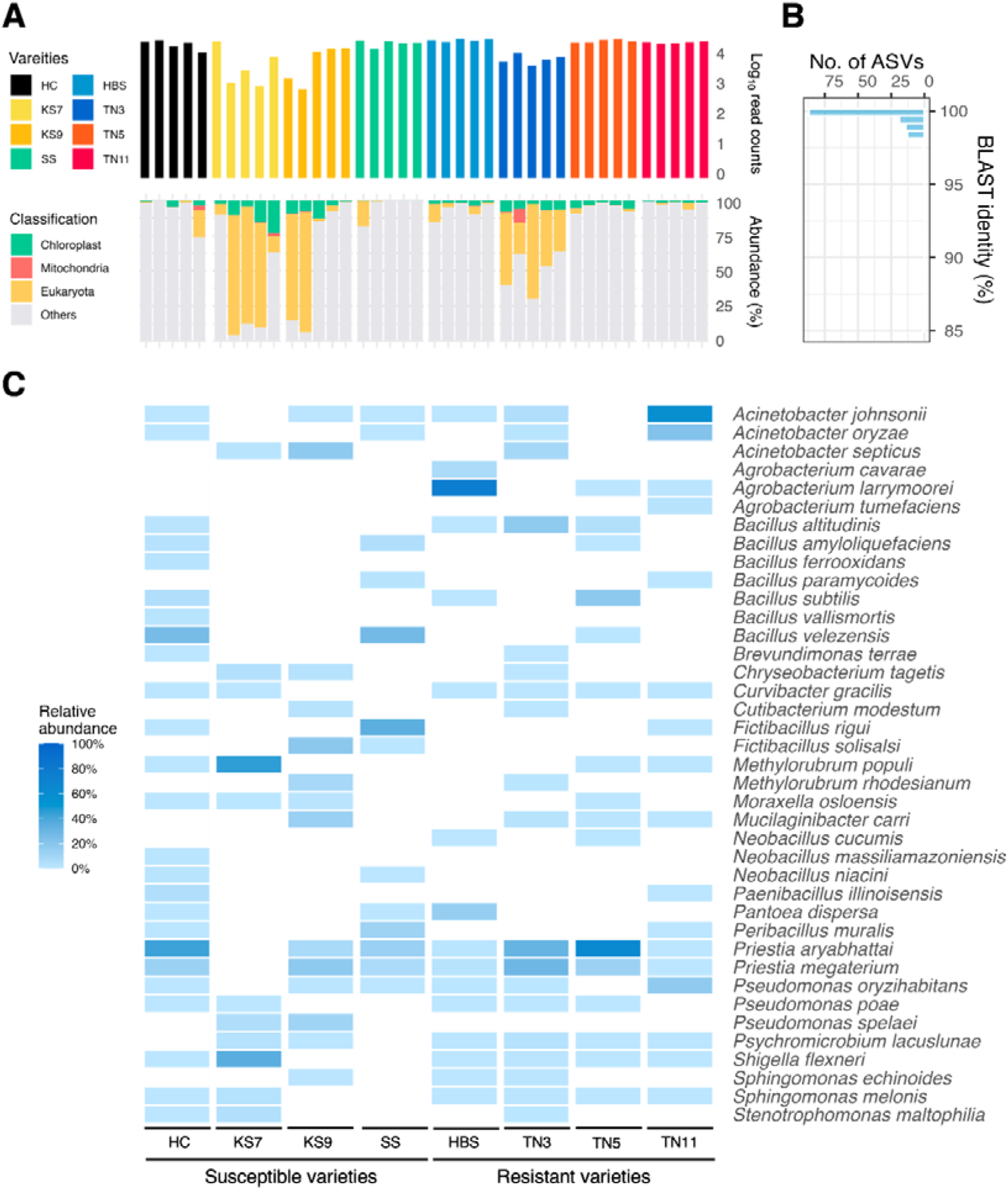
PacBio 16S rDNA full-length analyses to identify seed-associated bacteria in the eight soybean varieties. **A** The upper bar plot shows the read counts of each sample after removing host contamination DNA. The lower stacked bar plot shows the proportion of the host-contaminated DNA of each sample. **B** The BLAST identity for ASVs compared to the reference taxa. **C** Heatmap of relative abundance for the bacterial species in each soybean variety.

The taxonomic profile unveiled four bacterial phyla associated with soybean seeds, with the predominant taxa being Bacillota (56.4%), Pseudomonadota (41.0%), Bacteroidota (1.9%), and Actinomycetota (0.7%). At the family level, Bacillaceae emerged as the most abundant family (55.7%), followed by Moraxellaceae (15%) and Rhizobiaceae (11.7%) (Table S5). At the species level, the 135 ASVs were attributed to 39 bacterial species. Two bacterial species were identified in seven varieties except for KS7. Six bacterial species were present in six varieties, and one bacterial species was detected in five varieties (Fig. 3C). Additionally, 25 bacterial species were found in fewer than four varieties. The results indicated a considerable variability of the bacterial composition across soybean varieties.

### **α**-diversity, **β**-diversity, and network analysis of the seed microbiome

Indeed, the β-diversity analysis supported that the seed microbiome was predominantly shaped by the soybean variety (Fig. 4A). PERMANOVA disclosed that the soybean variety contributed to 46.6% of the total variance in microbial composition (Table S6). The hierarchical cluster analysis, based on the presence or absence of 39 bacteria, revealed clear groupings according to soybean varieties (Fig. S3A). Furthermore, richness (*p* = 0.002), Shannon diversity (*p* = 0.005), and Pielou’s evenness (*p* = 0.002) exhibited significant differences among soybean varieties (Fig. S3B). On the other hand, the seed rot resistance explained 7.1% of the total variance, and α-diversity analysis between the resistant and susceptible varieties showed no differences across all five indices (Fig. 4B). Network analyses indicated greater dissimilarity between the soybean varieties than between the seed rot resistance (Table S7). The network topology also showed these ASVs forming discrete modules with sparse interconnectivity between modules for each soybean variety (Fig. 4C). Additionally, the hub species were distinct among varieties (Table S8), suggesting that inter-microbial relationships may not directly contribute to the seed rot resistance.

**Fig. 4.**
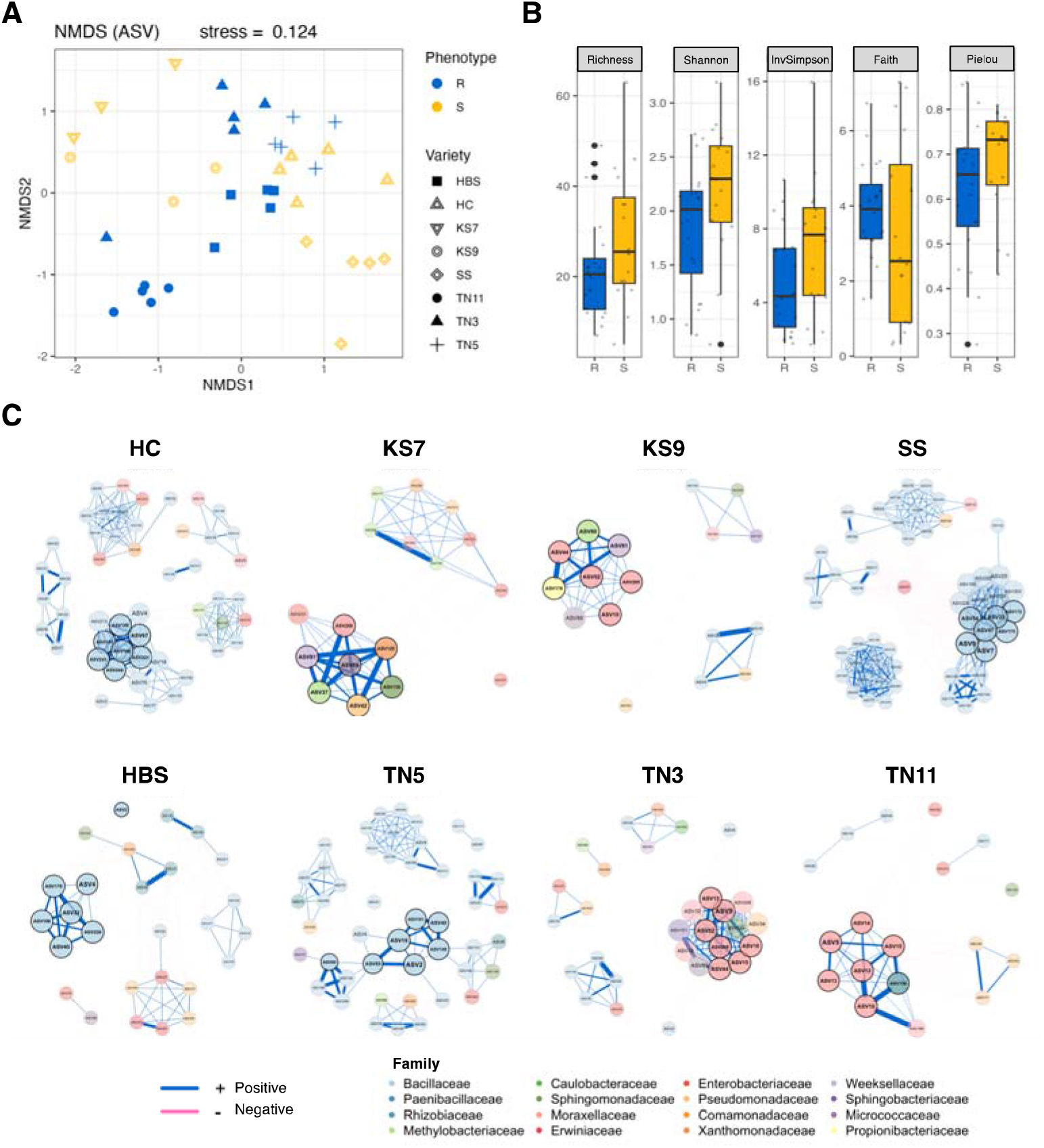
Bacterial diversity and network analysis of the eight soybean varieties. **A** The NMDS plot based on the Bray-Curtis distance. The colors represent the seed rot resistance or susceptibility, and the shapes represent the soybean varieties. **B** The α-diversity indices (richness, Shannon index, inverse Simpson index, Faith’s phylogenetic index, and Pielou’s evenness index) between the resistant and susceptible varieties. No significant difference was found by the *t*-test at α = 0.05. **C** The co-occurrence networks of the seed-associated bacteria in the eight soybean cultivars. Network inference was performed at the ASV level, and each node corresponded to a unique ASV. The size of a node depicts the eigenvector centrality, while the colors represent the bacterial family. The hub nodes are represented by the bold circles. The blue edges indicate for correlations above 0.7 and the red edges for correlations below -0.7. The thickness of each edge indicates the strength of correlation.

Instead, differential abundance analysis identified 14 ASVs being more abundant in the resistant varieties (Fig. 5). These ASVs were assigned to seven bacterial species, and the most enriched species included *Agrobacterium cavarae* ASV35 and ASV38, *Agrobacterium larrymoorei* ASV3 and ASV6, and *B. altitudinis* ASV20 and ASV50 (Table S5). Additionally, in the resistant soybean varieties, *Acinetobacter johnsonii* ASV5 and ASV13 were identified as hub species in TN3, *Priestia aryabhattai* ASV55 in TN5, and *Acinetobacter johnsonii* ASV12 along with *Acinetobacter oryzae* ASV14 in TN11 (Fig. 4C; Table S8).

**Fig. 5.**
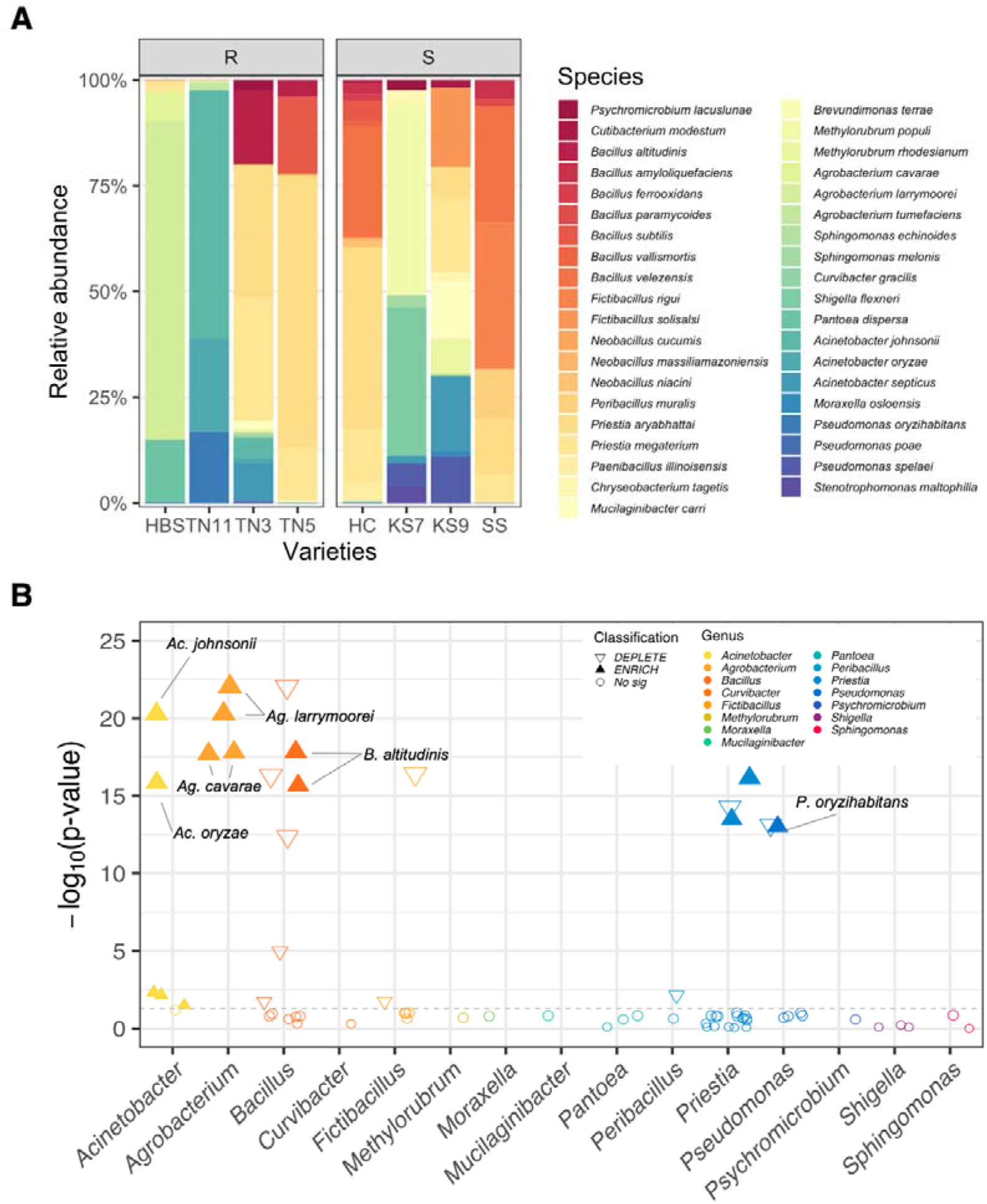
Taxonomic profile of the seed-associated bacteria between the resistant and susceptible varieties. **A** The proportion of the seed-associated bacteria at the species level. The reads counts were normalized by the medium of sequencing depth and covert to the percentage of the composition. **B** The Manhattan plots showing the ASVs, which are represented by circles or triangles. While the circles are insignificant ASVs in the differential abundance analysis, the triangles are ASVs significantly enriched (filled) or depleted (empty) in the resistant varieties. The triangle size indicates the log_2_ fold change of the ASV.

### Identification of *Bacillus altitudini*s to inhibit fungal pathogens

A total of 300 bacterial isolates were gathered from the seeds of eight soybean varieties, and 93 isolates with distinct colony morphology were selected for the *in vitro* antagonistic assay against *C. ilicicola*. The results disclosed that 29 bacterial isolates inhibited at least 20% of the mycelial growth of *C. ilicicola* (Table S9). Among these isolates, one from TN3 and one from TN5 displayed a clear inhibition zone. Molecular identification and phylogenetic analysis of 16S rRNA and *gyrB* gene sequences revealed that both isolates belonged to *B. altitudinis* (Fig. S4).

Remarkably, these two bacterial isolates (TN3S3 and TN5S8) exhibited antagonistic activity against other soybean soil-borne pathogens, inhibiting the mycelial growth over 30% for *A. rolfsii*, *M. phaseolina*, *R. solani*, and *S. sclerotiorum*, and over 20% for *F. oxysporum* (Fig. 6). Using TN5S8 in the subsequent experiments, the results demonstrated that upon re-inoculation to antibiotics-treated soybean seeds (Fig. 7A, B), TN5S8 significantly mitigated seed mortality and cotyledon rot caused by *C. ilicicola* across all resistant varieties and the susceptible variety KS9 (Fig. 7C, D). These findings strongly suggest that *B. altitudinis* T5S8 played a pivotal role in conferring the seed rot resistance against *C. ilicicola*.

**Fig. 6.**
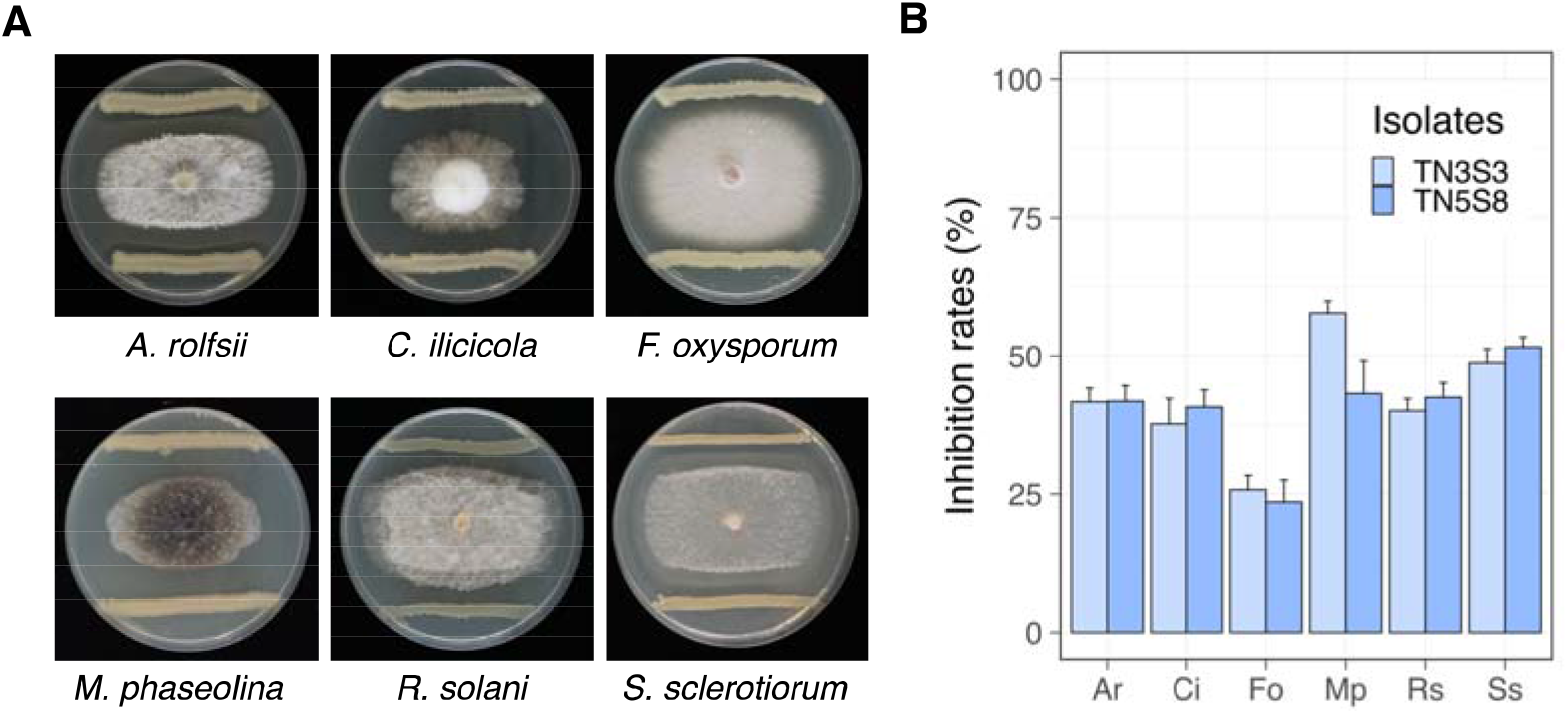
*In vitro* assay of *B. altitudinis* TN5S8 and TN3S3 against six soil-borne fungal pathogens **A** Representative image of the dual culture plates using TN5S8 to antagonize six fungal pathogens. **B** The inhibition rates. The values are the means[±[standard deviation. Ar: *Athelia rolfsii*, Ci: *Calonectria ilicicola*, Fo: *Fusarium oxysporum*, Mp: *Macrophomia phaseolina*, Rs: *Rhizoctonia solani*, Ss: *Sclerotinia sclerotiorum*.

**Fig. 7.**
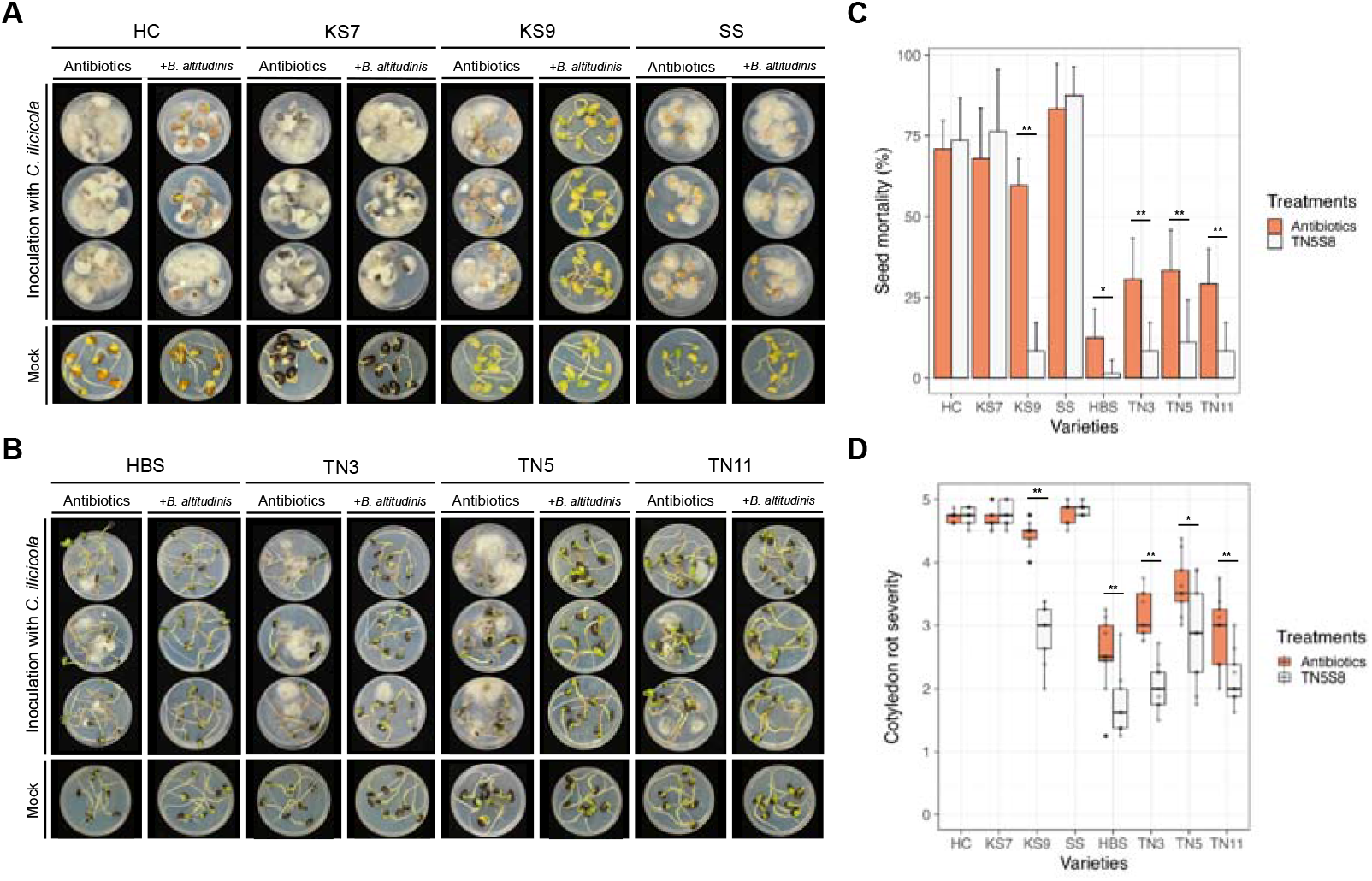
Seed rot assay using the antibiotics-treated seeds with or without the inoculation of *B. altitudinis* TN5S8. **A** The four susceptible soybean varieties with three showed no difference between the antibiotic treatment and the application of TN5S8. KS9 is the only variety being rescued by the application of TN5S8. **B** All four resistant soybean varieties showed significant reduction in the seed mortality and the cotyledon rot with the application of TN5S8. **C** Seed mortality. **D** Cotyledon rot severity. The values are presented as the means ± standard deviation. The asterisks indicate significance based on the *t*-test (*: *p* < 0.01, **: *p* < 0.001).

### Colonization of *Bacillus altitudinis* TN5S8 on soybean is a prerequisite to gain the seed rot resistance in compatible soybean varieties

However, the introduction of TN5S8 did not alleviate the seed rot susceptibility for three susceptible varieties—HC, KS7, and SS. We postulated that the compatibility between TN5S8 and soybean varieties might be a pivotal factor in gaining the seed rot resistance. Therefore, we quantified the colonization of TN5S8 on various soybean varieties and observed a significantly higher recovery of TN5S8 in all resistant varieties and the susceptible variety KS9 (Fig. 8A). Notably, the highest recovery of TN5S8 was observed from the original isolation variety TN5, indicating a preference of TN5S8 for specific soybean varieties. Additionally, using qPCR to detect the presence of TN5S8 on the TN5 seedlings, TN5S8 was found to persist on the apical shoot and first node of soybean seedlings until 21 dpi, but it was not detected on the roots after 9 dpi (Fig. 8B). These results collectively suggested that TN5S8 may initially colonize the seeds and subsequently migrate to the shoots.

**Fig. 8.**
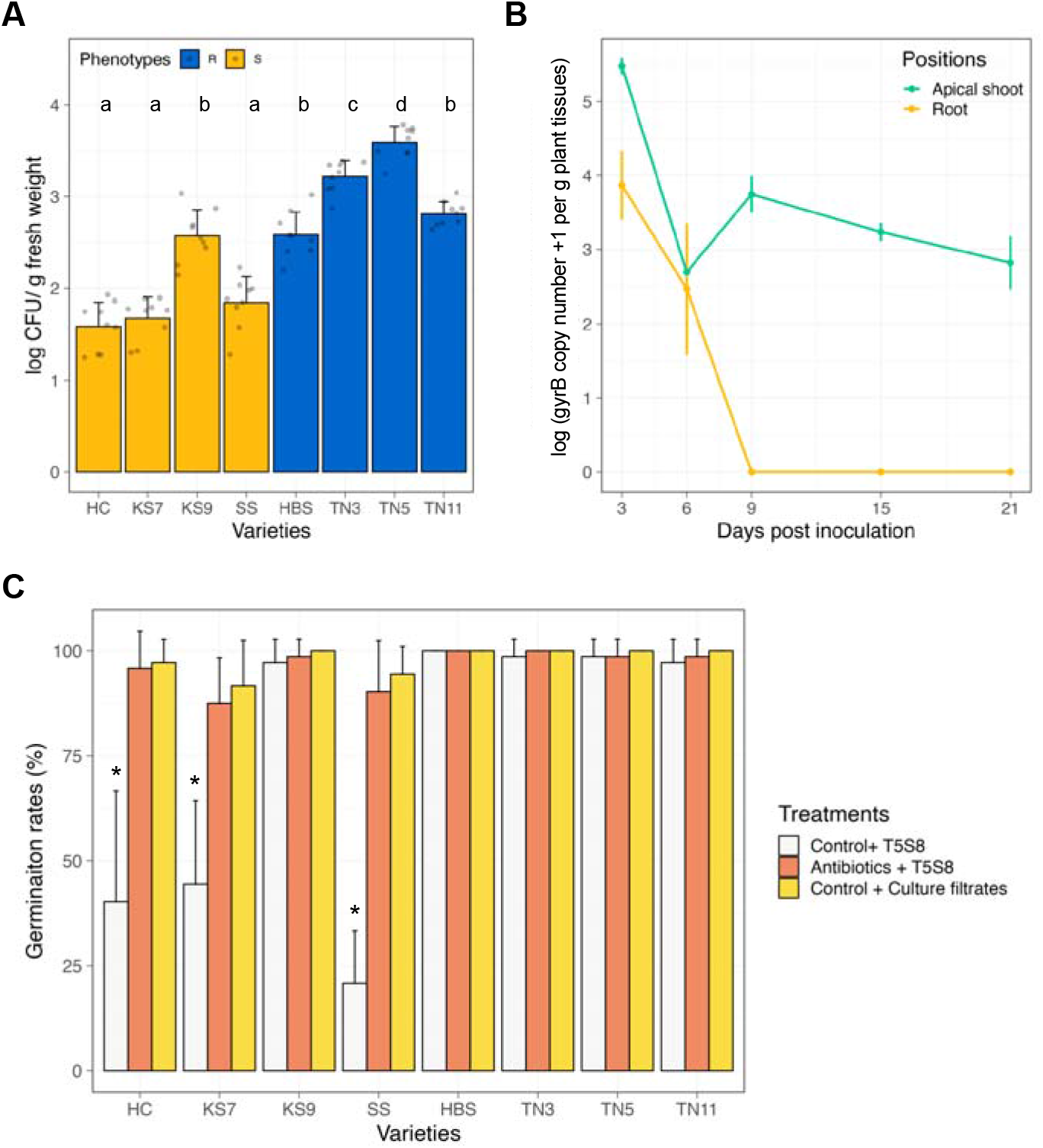
Compatibility of *B. altitudinis* TN5S8 on the seeds of eight soybean cultivars. **A** The recovery of TN5S8 abundance on the antibiotic-disinfected seeds at 5 dpi. The colors indicate the seed rot resistance or susceptibility. The values are means[±[standard deviation. ANOVA and the Tukey’s HSD test were used to determine the significance at α = 0.05. **B** Quantitative PCR (qPCR) of *B. altitudinis* on the soybean apical shoots or roots at different timepoints. **C** The germination rates of soybean seeds. The white bars indicate the application of TN5S8 on the surfaced-sterilized seeds without antibiotic treatment. The orange bars indicate the application of TN5S8 on the surfaced-sterilized seeds with antibiotic treatment. The yellow bars indicate the application of TN5S8 culture filtrate, instead of bacteria, on the surfaced-sterilized seeds without antibiotic treatment. The values are means[±[standard deviation. The Kruskal-Wallis and the Dunn’s test were used to determine significant difference at α = 0.05.

Interestingly, TN5S8 significantly impeded the seed germination of soybean varieties HC, KS7, and SS (Fig. 8C), indicating lower compatibility between TN5S8 and these three varieties (Fig. 8A). The non-germinated seeds of HC, KS7, and SS exhibited not only a dark hue but also soft rot. Importantly, the germination reduction and associated symptoms were not induced by the cell-free culture filtrate of TN5S8. These findings underscored the importance of compatibility between TN5S8 and soybean varieties to confer the seed rot resistance.

## Discussion

Plant genotypes have been recognized as the major factor contributing to disease resistance, not only for soybean [56] but also for most important crops [57, 58]. However, some studies have observed that fungal infection on different tissues such as seed, root, node, or leaf of the same plant genotype could result in different levels of resistance [59, 60]. For example, a study of soybean resistance to *Pythium* revealed the phenotypic correlation between seed rot and root rot was between 0.1 and 0.17 [59]. Another study on the pea resistance to *S. sclerotiorum* reported the phenotypic correlation of nodal resistance and leaf resistance was only 0.19. Recently, it has been known that disease resistance of plants can also be governed by the host-associated microbiome [61–64] Specifically for the seed-associated bacteria and disease resistance, distinct bacterial compositions in the seed of different oilseed rape cultivars were correlated with varying resistance levels to Verticillium wilt and *Plasmodiophora brassicae* [13, 65]. In another study, a seed endophytic bacterium *Sphingomonas melonis*, which can be vertically transmitted to the next generation of seeds, was found to secrete anthranilic acid to confer rice seedlings resistance against *Burkholderia plantarii* [5]. Similarly, *Bacillus velezensis* isolated from maize seeds and *Bacillus subtilis* found in millet seeds were shown to protect seedlings from *Fusarium* infection [22, 23]. However, these studies only focused on the roles of seed-associated bacteria in conferring seedling resistance, leaving it unclear whether the seed-associated bacteria can protect seeds and whether it may also involve in the discrete resistance, differing in plant tissues of the same genotype.

In assessing 16 soybean varieties, this study identified a discrete resistance, present only for seed rot, but not for root rot. The discrete resistance between seed rot and root rot suggested that the underlying mechanisms might involve factors other than plant genotypes, leading us to the hypothesis that the seed-associated bacteria may confer to the seed rot resistance, and it may not be carried to the roots.

Based on the experiments using the antibiotics-treated seeds, and the results confirmed that the seed-associated bacteria were involved in the seed rot resistance. As previously reported that the α-diversity or co-occurrence network properties between the resistant and susceptible plants were different [65, 66] and may protect the resistant plants from pathogen by increasing niche overlap, competing for resource consumption, or intensifying antagonistic interactions [67], this study applied 16s rDNA full-length sequencing and microbiome analyses to compare the seed microbiome between the resistant and susceptible soybean varieties. Nevertheless, there was no significant differences in the α-diversity or co-occurrence networks. Instead, a relatively low bacterial diversity, with an average of merely 16 bacterial species, was identified in the seeds of one variety, similar to other seed microbiome studies [34, 36]. Consistently, the differential abundance analysis discovered bacterial species that were significantly enriched in the resistant varieties (Table S6), suggesting that a certain group of seed-associated bacteria, rather than a community, may contribute to the seed rot resistance.

There were 14 ASVs enriched in the resistant soybean varieties, including species such as *A. johnsonii*, *A. oryzae*, *A. larrymoorei*, *A. cavarae*, *B. altitudinis,* and *P. oryzihabitans. A. johnsonii* [68] has been previously isolated from soybeans, exhibiting antagonistic capabilities against soil-borne pathogens. In addition, *A. larrymoorei* and *P. oryzihabitans* have been reported as soybean endophytic bacteria, showing potential in the nitrogen fixation and phosphate solubilization. [69, 70]. Moreover, some *P. oryzihabitans* strains have antagonistic capability against pathogens such as *Acidovorax citrulli* in cucurbits and *Pythium* in cotton [71, 72]. While the literature suggested that these bacteria may play roles in the seed rot resistance, our culture-dependent isolation resulted in another enriched ASV, which was identified as *B. altitudinis*. In our experiments, loss-of-function evidence through the antibiotic treatment and the gain-of-function evidence through the application of *B. altitudinis* to soybean seeds confirmed the contribution of *B. altitudinis* in the seed rot resistance.

*B. altitudinis* was first isolated from extreme UV-stressed air samples collected in the stratosphere [73]. It has been identified as an endophyte in various plants, including soybean [74] and others [75, 76, 77, 78, 79, 80, 81]. Several strains of *B. altitudinis* have shown biocontrol capabilities, such as cotton Verticillium wilt [75], grape downy mildew [78], kiwi fruit root-knot nematodes [82], soybean Phytophthora damping-off [74] and sweet potato black rot [80]. It has been shown that *B. altitudinis* can inhibit plant pathogens by producing antimicrobial lipopeptides lichenysin [78, 83] and inducing plant defense responses [74, 82]. Moreover, genome analysis also identified genes required to synthesize fengycin and bacilysin, and these compounds may be associated with the antagonistic ability [84]. Recently, *B. altitudinis* has been suggested to have an open pangenome, and 42.7% genes were characterized as accessory genes (2,349 genes). The phenomenon indicated that *B. altitudinis* tends to acquire new genes to enhance its antagonistic capability and ecological competitiveness [78].

We observed that the success of seed rot resistance in soybeans depends on the colonization of TN5S8 on the compatible soybean varieties. The relationship between bacterial population and disease suppression echoes previous findings on the biocontrol efficacy of *Pseudomonas fluorescens* was proportional to their population density [85], and the effective threshold ranges from 10^5^ to 10^6^ cells per gram of root against wheat take-all decline disease [86]. Similarly, the suppression of other Pythium root diseases in sugar beets also depended on the population density of the *Pseudomonas* [87, 88]. In rice, the abundance of *Sphingomonas* was observed to be lower in plants susceptible to seedling blight disease [5]. Specifically for the cases within the *Bacillus* genus, colonization and formation of biofilm on the phyllosphere or root surface is critical for the success of biocontrol [89]. On tomato, *Bacillus* strains with less colonization ability on the phyllosphere showed a reduced biocontrol ability against *Botrytis cinerea* [90]. Application of plant extracts such as pectin can enhance the population density of *Bacillus amyloliquefaciens* on tobacco roots and increase the biocontrol efficacy to tobacco bacterial wilt [91]. Mutation of *B. amyloliquefaciens abrB* gene, which is a negative transcription regulator of chemotaxis and biofilm formation, increased colonization and biocontrol capability against the cucumber Fusarium wilt [92]. Moreover, the colonization of *B. subtilis* surfactin deletion mutant reduced 4 to 10-fold on the melon roots and leaves, which ended up losing the biocontrol efficacy [93]. However, it has not be reported whether the colonization of *Bacillus* can affect seed resistance, and our finding provided the first evidence that the colonization of *B. altitudinis* on soybean seeds is an important factor to confer the seed rot resistance.

It has been proposed that not all seed microbiota can migrate from seeds to seedlings [94, 95]. A recent study on oak showed that 63% of fungal taxa and 45% of bacterial taxa on the seeds can be transmitted to the seedlings [15]. Another study on tomato demonstrated that some seed-associated microbes such as *Bacillus aryabhattai*, *Bacillus nakamurai*, *Ralstonia pickettii*, and *Stenotrophomonas maltophilia* could persist from seeds to seedlings for at least two generations [11]. However, even though the seed-associated bacteria can be transmitted to seedlings, their colonization on the shoots or roots may be different. A study on soybeans in an axenic environment demonstrated that the seed-transmitted bacterial ASVs dominant in the shoots can be rare or absent in the roots [34]. This report aligns with our observation on TN5S8, which was detected on the apical shoots for at least 21 days, but it could not be detected on the root 9 days. This time-course experiment suggested that TN5S8 may migrate to the aboveground parts of soybean during plant growth, but the absence of TN5S8 on the roots resulted in no protection in the root rot assays.

Bacteria must tackle plant immune responses to successfully establish colonization [96]. Therefore, it is anticipated that the colonization of commensal bacteria could vary among different plant genotypes or tissues. In this study, we show that the community of soybean seed microbiome was predominantly variety-dependent, and the seed-associated bacteria such as *B. altitudinis* TN5S8 may provide antagonistic capability to other fungal pathogens. Moreover, this study pointed out that *B. altitudinis* TN5S8 confers only seed rot resistance in certain soybean varieties based on its colonization compatibility. The results highlight for future application of the seed-associated bacteria in disease management to consider not only the antagonistic capability, but also the colonization compatibility on the plant genotypes and tissues.

## Data availability

Raw sequencing sequences have been deposited in the NCBI Sequence Read Archive (SRA) under the BioProject PRJNA1019790.

## Supplementary Figures

**Supplementary Fig. S1.**
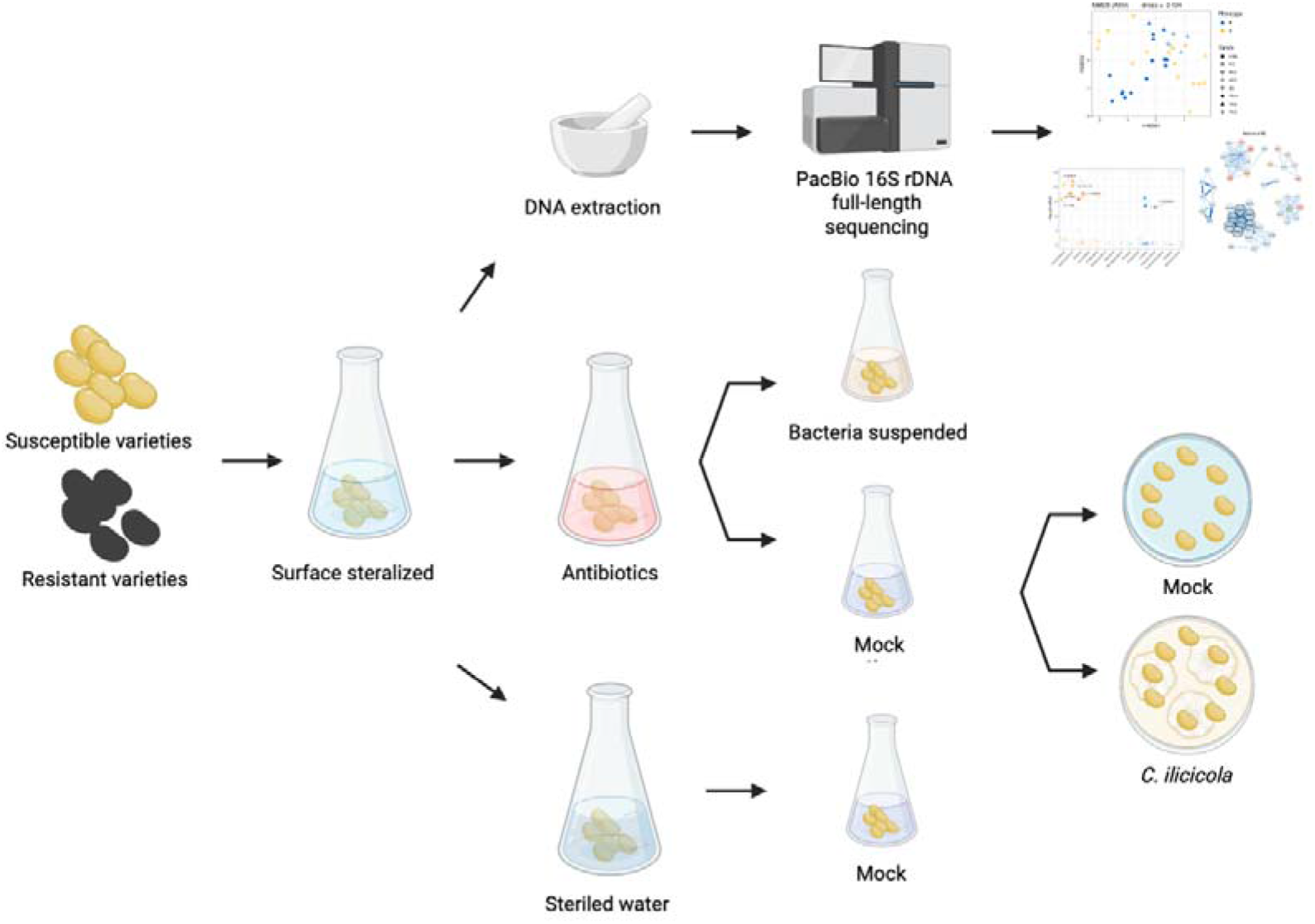
Schematic workflow of the antibiotics-treated soybean seeds to study the potential of the seed-associated bacteria contributing to the seed rot resistance.

**Supplementary Fig. S2.**
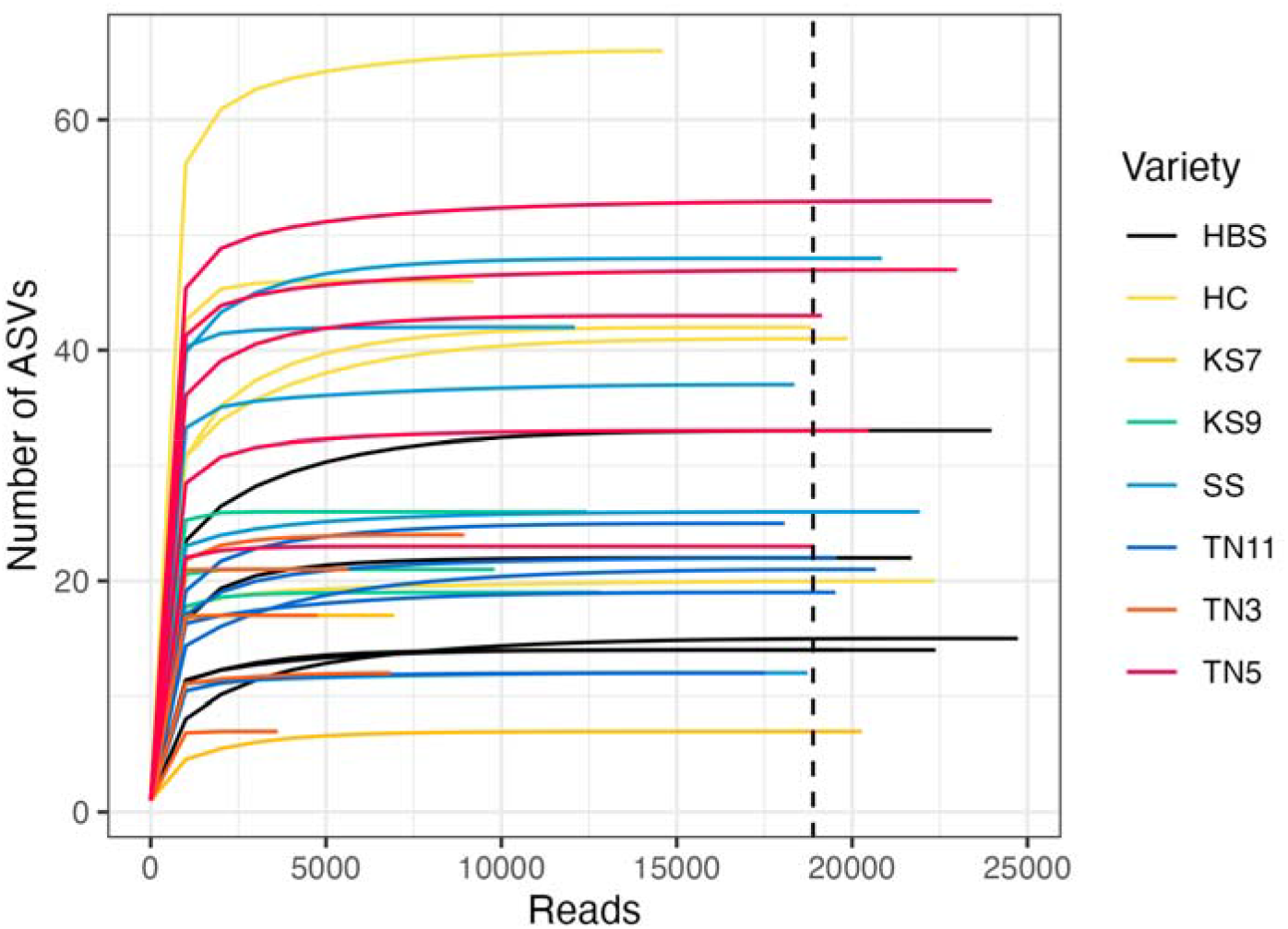
Rarefaction curves of all samples. The dashed line indicates the median read depth.

**Supplementary Fig. S3.**
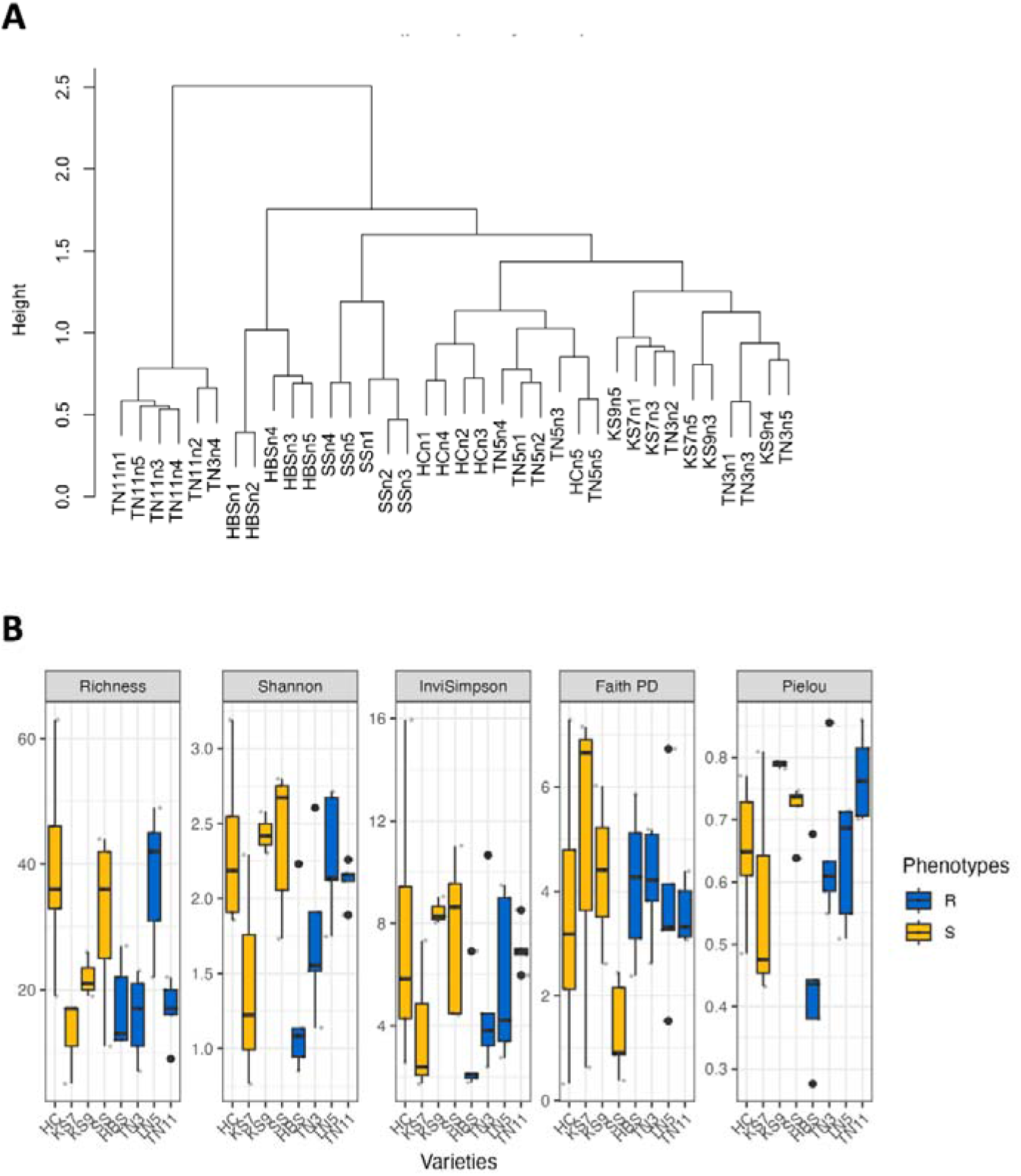
Cluster and α-diversity analysis of the seed-associated bacteria in the eight soybean varieties. A Cluster analysis by the beta flexible method (with α = 0.625) using the Sørensen distance. B α-diversity. The colors indicated the seed rot resistance (blue) or susceptible (yellow).

**Supplementary Fig. S4.**
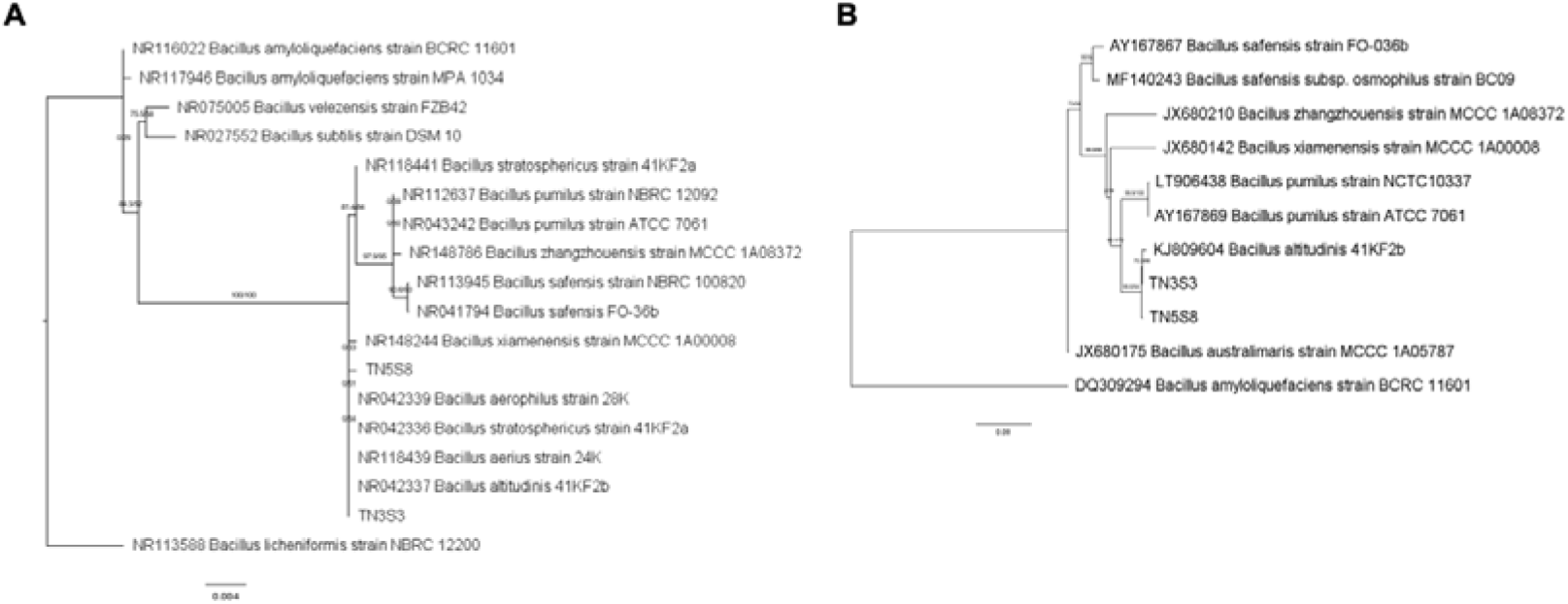
Identification of species identities for TN5S8 and TN3S3 using the maximum likelihood (ML) phylogenetic analysis. A ML phylogenetic tree based on the 16S rRNA gene. B ML phylogenetic tree based on the *gyrB* gene. Numbers indicated the bootstrap values and the Shimodaira–Hasegawa approximate likelihood ratio test (SH-aLRT) values.

## Notes

### Competing Interest Statement

The authors have declared no competing interest.

